# Coronatine is More Potent than Jasmonates in Regulating Arabidopsis Circadian Clock

**DOI:** 10.1101/2020.06.15.152439

**Authors:** Min Gao, Chong Zhang, Hua Lu

## Abstract

Recent studies establish a crucial role of the circadian clock in regulating plant defense against pathogens. Whether pathogens modulate host circadian clock as a potential strategy to suppress host innate immunity is not well understood. Coronatine is a toxin produced by the bacterial pathogen *Pseudomonas syringae* that is known to counteract Arabidopsis defense through mimicking defense signaling molecules, jasmonates (JAs). We report here that COR preferentially suppresses expression of clock-related genes in high throughput gene expression studies, compared with the plant-derived JA molecule methyl jasmonate (MJ). COR treatment dampens the amplitude and lengthens the period of all four reporters tested while MJ and another JA agonist JA-isoleucine (JA-Ile) only affect some reporters. COR, MJ, and JA-Ile act through the canonical JA receptor COI1 in clock regulation. These data support a stronger role of the pathogen-derived molecule COR than plant-derived JA molecules in regulating Arabidopsis clock. Further study shall reveal mechanisms underlying COR regulation of host circadian clock.

## Introduction

Plants have evolved sophisticated mechanisms to respond to daily attacks of pathogens and pests of different lifestyles. Increasing evidence has established that the circadian clock is an integral part of the plant innate immune system. In addition to being crucial for plant growth and development, the circadian clock regulates multiple layers of defense responses, including stomatal opening and closure, pathogen recognition, and defense signal activation [1, 2]. Whether the circadian clock is modulated by pathogens and pests as a potential strategy to circumvent host defense has not been well understood.

Jasmonates (JAs) are lipid-derived molecules that are important for defense signaling. Recent studies demonstrate a circadian regulation of JA signaling. The JA level and expression of some key JA biosynthetic and signaling genes oscillate in a day [3, 4]. Two Arabidopsis clock genes, LUX [5] and TIC [4] were shown to regulate clock output to affect JA signaling. In turn, JA signal activation dampens the amplitude and lengthens the period of some clock reporters [5], suggesting a reciprocal regulation of the circadian clock by JA signaling. Such a reciprocation relationship is also found between the circadian clock and other biological processes, including nutrient uptake and signaling mediated by salicylic acid, reactive oxygen species, and phytohormones [6–9], and it suggests an adaptive nature of plants to coordinate limited resources for growth, development, and environmental stimuli.

While being important for plant defense, JA signaling succumbs to pathogen interference. The bacterium pathogen, *Pseudomonas syringae*, is known to employ several mechanisms to manipulate host JA signaling, for instance, using the phytotoxin compound coronatine (COR) to mimic JA or using effector proteins to interfere with JA signaling [10–13]. COR structurally mimics JA-isoleucine (JA-Ile), a bioactive form of JA. Both COR and JA-Ile bind to the JA receptor, CORONATINE INSENSITIVE 1 (COI1), to activate JA signaling. COR is also known to regulate other defense pathways independently of its promotion of JA signaling [14]. Thus, COR and JA-Ile have overlapping and also distinct function in regulating biological processes. We recently showed that activation of JA signaling, using JA-Ile or another JA analog methyl jasmonate (MJ), reciprocally regulates clock activity [5]. Whether pathogen-derived COR could modulate clock activity has not been tested prior to this study. We report here that compared with plant-derived JA molecules, COR shows a stronger regulation of the circadian clock in Arabidopsis.

## Results

COR is a phytotoxin produced by pathovars of *P. syringae* and is important for the pathogenesis of the bacteria. The lack of COR makes *P. syringae* less virulent under a diurnal light and dark (LD) cycle [15, 16]. In continuous light (LL), a free-running condition often used to test clock activity, we found that compared with *P. syringae* strain DC3000, the isogenic *P. syringae* strain DC3118 that does not produce COR, grew much less and induced less chlorosis and lesion in the infected Arabidopsis leaves (Figure S1).

One way that COR promotes bacterial virulence is through interfering with JA signaling. A number of studies demonstrated crosstalk between JA signaling and the circadian clock [3–5, 17]. We are interested in elucidating in this report whether *P. syringae*-derived COR can regulate plant circadian clock. Toward this goal, we first compared expression of a set of circadian genes (Table S1), using two sets of time-series RNA-seq data from samples treated with 100 μM MJ or 5 μM COR [18, 19]. Heatmap analysis showed that COR induced more changes in expression of the circadian genes than MJ (Figure 1A and 1B). We estimated the relative number of affected gene expression by normalizing the total number of affected expression events with the number of time points. While the number of induced circadian genes was similar with the two treatments, COR showed a stronger suppression of circadian genes than MJ, including some core clock genes (Figure 1C and S2). We performed a similar analysis with a set of defense genes (Table S2). Interestingly, MJ and COR showed less difference in affecting defense gene expression (Figure 1C and S3).

**Figure 1.**
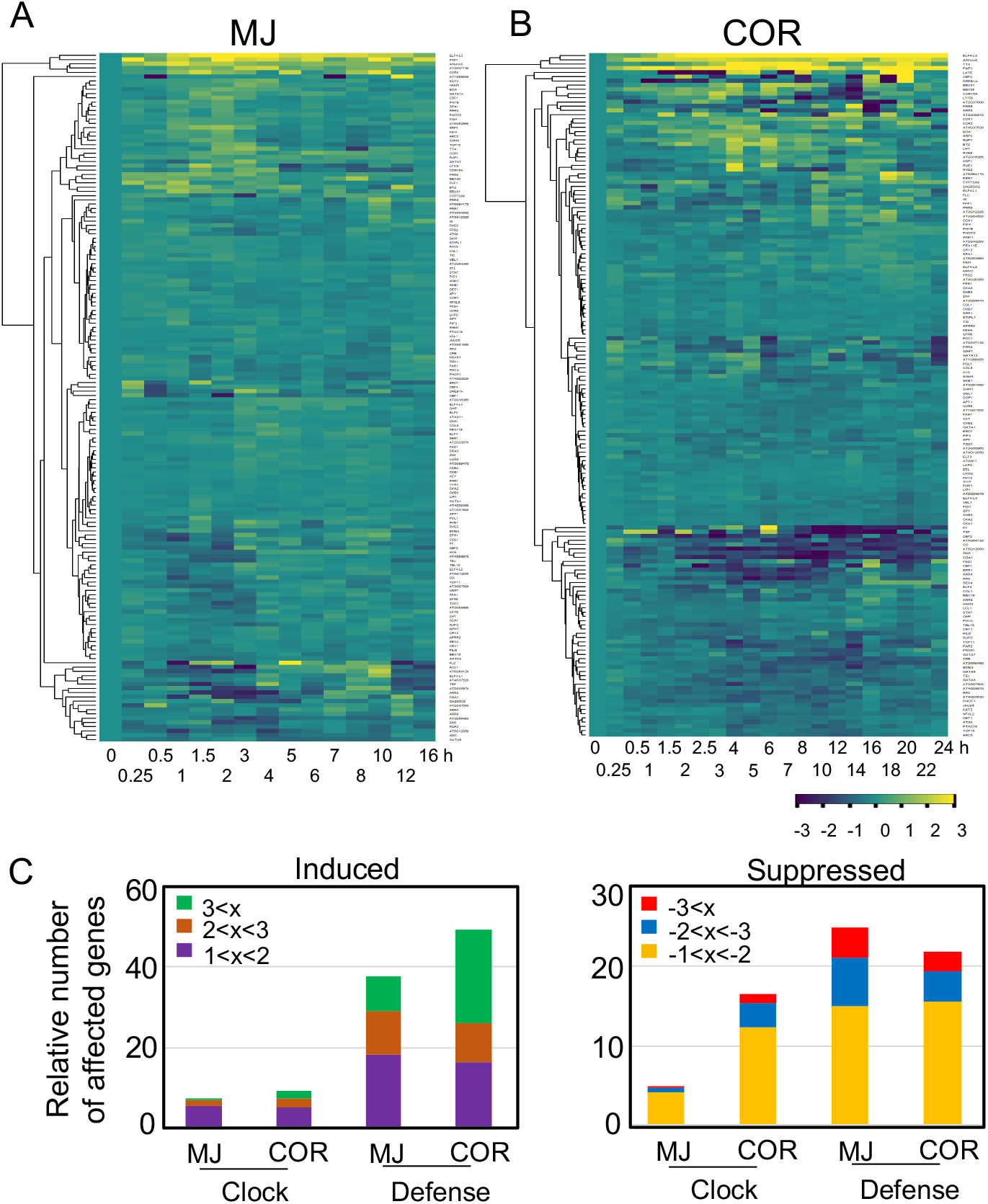
COR exerts stronger suppression on expression of **circadian** genes than MJ. (A) Heatmap analysis of expression of circadian genes in MJ-treated samples. (B) Heatmap analysis of expression of circadian genes in COR-treated samples. For (A) and (B), Log_2_ transformed fold change of gene expression (100 μM MJ or 5 μM COR treatment vs. mock treatment) was used to generate the heatmaps with the heatmap.2 function in R package gplots. For MJ treatment, the mock solution contained 0.015% (v/v) Silwet L77 and 0.1% ethanol [19]. For COR treatment, water was used as a mock treatment [18]. (C) Relative number of genes affected by MJ or COR. Expression of each gene at a time point was considered as one gene expression event. The total number of defense or clock gene expression events in each category was normalized by the total number of time points.

While these analyses suggest that COR exerts a stronger suppression of circadian genes than MJ, we recognized that the two RNA-seq experiments were conducted in two different laboratories that used plants grown in different conditions [18, 19]. In addition, both experiments were conducted under diurnal cycles. To reconcile these differences and examine the role of COR in clock regulation, we grew seedlings for 7 d under a 12 h light and 12 h dark (LD) cycle for entrainment. We then transferred the seedlings to continuous light (LL) for 1 d and treated them with MJ (100μM) or COR (10μM) for gene expression analysis. qRT-PCR results showed that both MJ and COR suppressed expression of selected core clock genes (Figure S4 and [5]), supporting that both MJ and COR regulate clock activity.

To further test the clock regulatory role of COR, we performed the luciferase (LUC) assay with plants expressing the *LUC* gene driven by promoters of different clock genes, including *CCA1*, *TOC1*, *PRR7*, and *GRP7*. Similar to MJ, we found that COR suppressed seedling growth in our clock assay condition (Figure S5 and [5]). After normalizing the LUC amplitude to relative leaf area of seedlings, we observed that COR dampened the amplitude and lengthened the period of all four reporters largely in a dosage dependent manner, regardless COR was applied at 25 or 37 h after light onset (subjective dawn or subjective dusk, respectively) (Figure 2). COR further induced a lagging phase with the *TOC1:LUC* and *GRP7:LUC* reporters (Figure 2D2 and 2D4), suggesting a higher sensitivity of these two reporters to COR than other reporters tested. To test if this clock regulatory role of COR requires intact JA signaling, we used the JA receptor mutant (*coi1-17*) expressing *CCA1:LUC* [5, 20]. COR did not affect seedling growth and rhythmicity of the *CCA1:LUC* reporter in *coi1-17* (Figure 2A5, 2B5, 2C5, 2D5, and S5A). Thus, these results support that the role of COR in regulating clock activity requires a functional JA receptor.

**Figure 2.**
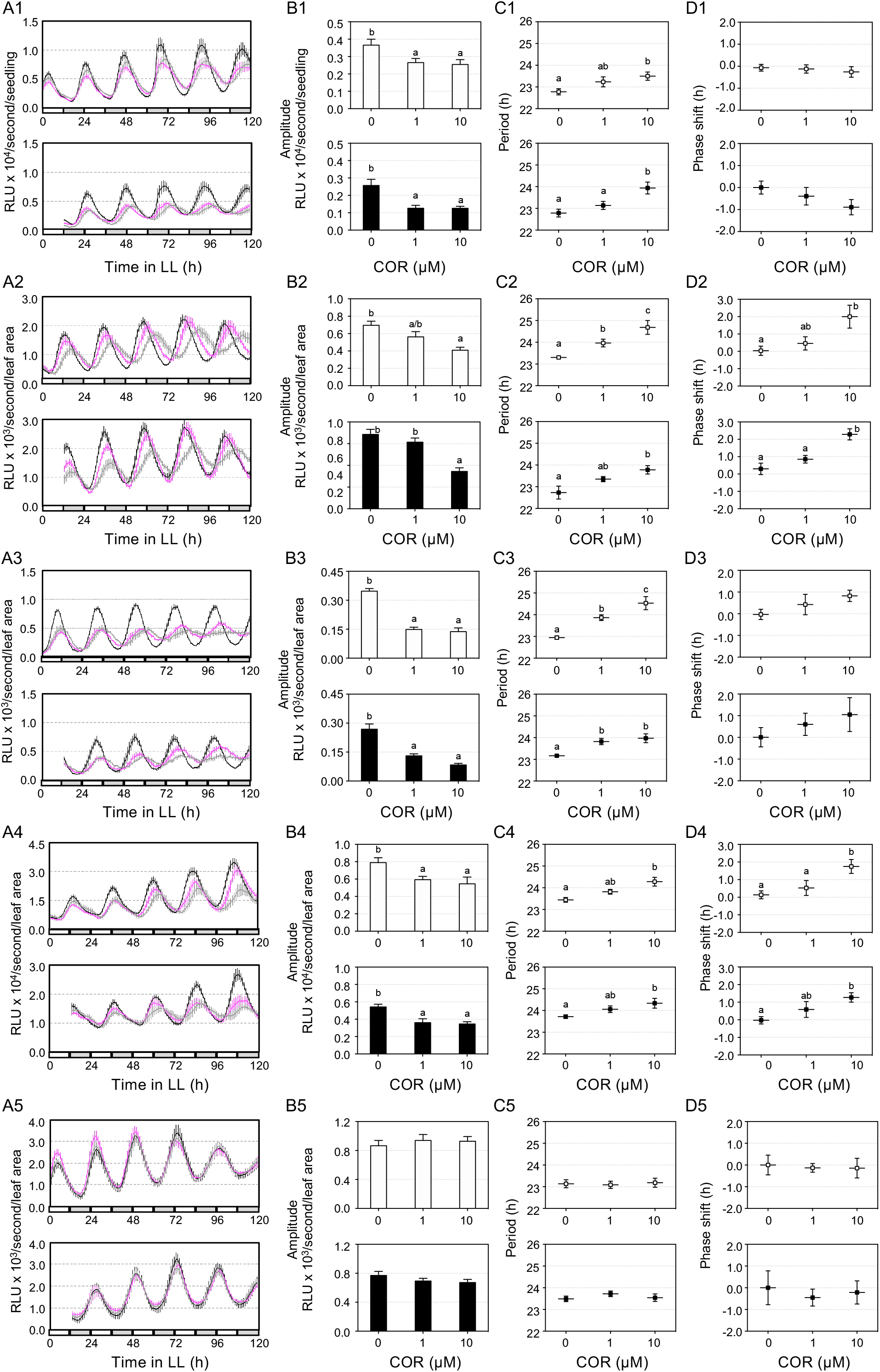
COR treatment affects clock activity. LD-entrained 5 d old seedlings were transferred to LL for 1 d and were treated with COR or water at 25 h (top of each panel) or 37 h (bottom of each panel). Luminescence was recorded at 1 h intervals for five days and analyzed for clock activity. A1-D1 Expression of *CCA1:LUC* in Col-0. A2-D2 Expression of *TOC1:LUC* in Col-0. A3-D3 Expression of *PRR7:LUC* in Col-0. A4-D4 Expression of *GRP7:LUC* in Col-0. A5-D5 Expression of *CCA1:LUC* in *coi-17*. A1-A5 Luminescence traces. RLU: Relative luminescence units. The color indicates COR concentration, black for 0, magenta for 1 μM, and gray for 10 μM. B1-B5 Normalized amplitude. The amplitude of the reporter was normalized to the relative leaf area shown in Figure S5. C1-C5 Period. D1-D5 Phase shift. Data represent mean ± SEM (n=12). Statistical analysis was performed by One-way ANOVA post-hoc Tukey HSD test. Different letters indicate significant difference among the samples (P<0.05). These experiments were repeated three times with similar results.

Like COR, MJ also affects seedling growth (Figure S5B). We previously showed that MJ only affects the amplitude but not the period and phase of the *CCA1:LUC* reporter [5]. We report here that three additional reporters (*TOC1:LUC*, *PRR7:LUC,* and *GRP7:LUC*) showed an amplitude dampening in the presence of MJ (Figure 3). The *PRR7:LUC* and *GRP7:LUC* reporters also displayed period lengthening, depending on MJ dosages. Furthermore, MJ induced phase lagging in *TOC1:LUC* and *GRP7:LUC*. Unlike COR, MJ did not affect the period of *CCA1:LUC* and *TOC1:LUC*. These results suggest a stronger effect of COR than MJ in regulating clock activity, at least for some clock genes. They also illustrate differential sensitivity of different clock reporters to COR, MJ, and JA-Ile.

**Figure 3.**
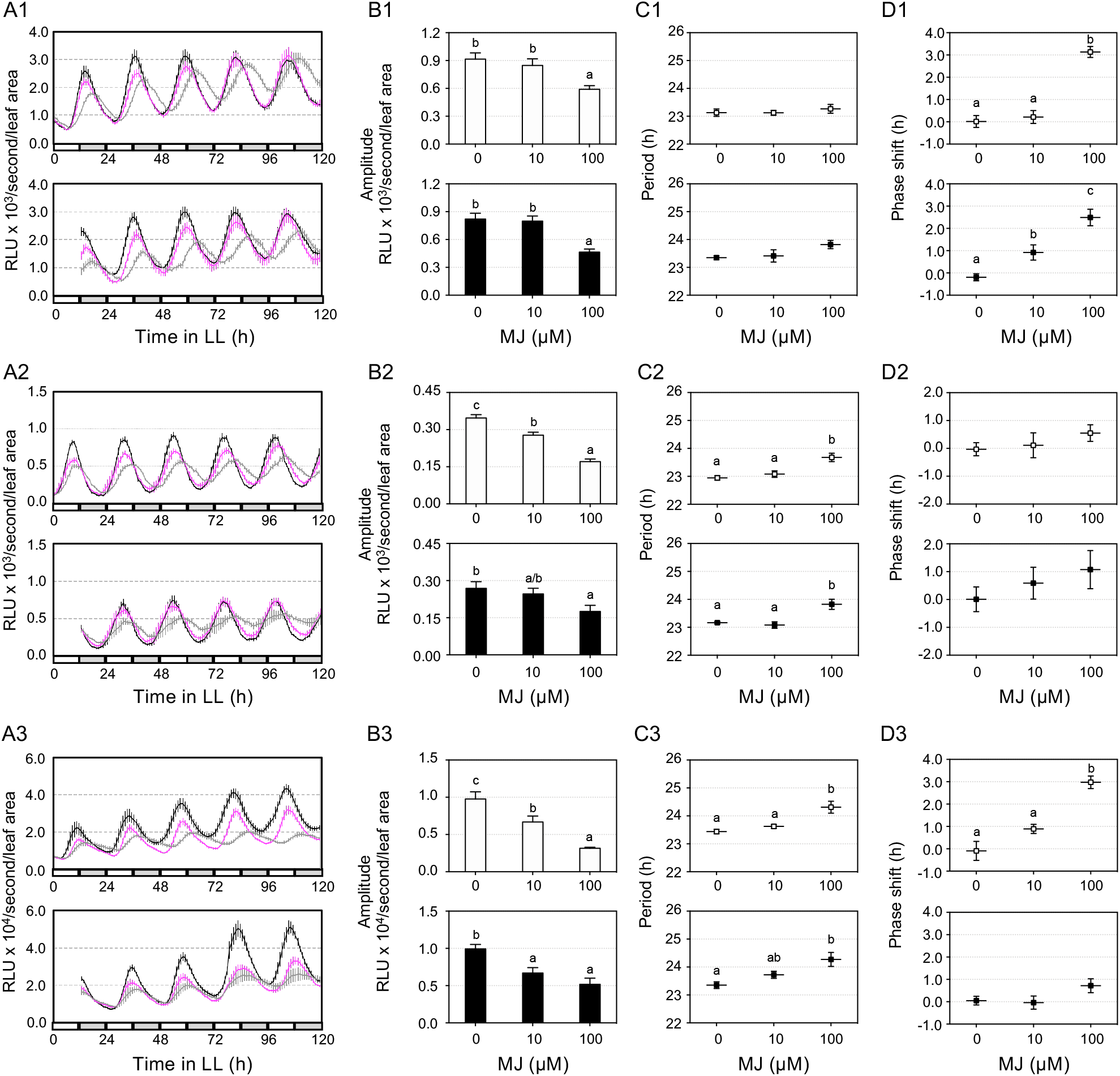
MJ treatment affects clock activity. LD-entrained 5 d old seedlings were transferred to LL for 1 d and were treated with MJ or water at 25 h (top of each panel) or 37 h (bottom of each panel). Luminescence was recorded at 1-h intervals for five days and analyzed for clock activity. A1-D1 Expression of *TOC1:LUC* in Col-0. A2-D2 Expression of *PRR7:LUC* in Col-0. A3-D3 Expression of *GRP7:LUC* in Col-0. A1-A3 Luminescence traces. RLU: Relative luminescence units. The color indicates MJ concentration, black for 0, magenta for 10 μM, and gray for 100 μM. B1-B3 Normalized amplitude. The amplitude of the reporter was normalized to the relative leaf area shown in Figure S5. C1-C3 Period. D1-D3 Phase shift. Data represent mean ± SEM (n=12). Statistical analysis was performed by One-way ANOVA post-hoc Tukey HSD test. Different letters indicate significant difference among the samples (P<0.05). These experiments were repeated three times with similar results.

We previously reported that another plant-derived jasmonate, JA-Ile, acts through COI1 to suppress the amplitude, lengthen the period, but not affect the phase of *CCA1:LUC* and *GRP7:LUC* reporters in Col-0 [5]. Seedling growth was not affected by JA-Ile (Figure S5C and [5]). We confirmed these results with the *PRR7:LUC* reporter (Figure S5C and S6). Interestingly, the *TOC1:LUC* reporter showed less sensitivity to JA-Ile than other reporters tested, only showing a dampened amplitude but no change in the period and the phase. These results support a stronger role of COR than JA-Ile in clock regulation and differential sensitivity of clock reporters to COR and plant JA derivatives.

## Discussion

Growing evidence indicates that pathogens can reprogram the circadian clock of the host. For instance, the bacterium *P. syringae*, the oomycete *Hyaloperonospora arabidopsidis,* and the fungus *Botrytis cinerea* were shown to manipulate the circadian clock of Arabidopsis [9, 21–23]. Even gut microbiota in the animal host are capable of reprograming their animal host clock [24, 25]. The key question remains how pathogens affect host clock activity and defense responses. Pathogens are known to secrete a vast range of molecules to interfere with host immunity. Studies just begin to reveal that some signals emanating from pathogens modulate the circadian clock of the host. Pathogen associated molecular patterns (PAMPs), including bacteria lipopolysaccharide (LPS) and flg22, were shown to affect the circadian system of animals and Arabidopsis, respectively [21, 26]. We report here that the *P. syringae*-produced toxin molecule COR exerts a stronger influence on Arabidopsis clock than some plant-derived JA molecules. Our conclusion is strongly supported by experimental evidence: 1) While MJ and COR similarly affected defense gene expression, COR showed a stronger suppression of circadian genes than MJ (Figure 1, S2, and S3); and 2) While COR, MJ, and JA-Ile dampened the amplitude of all four clock reporters in a dosage dependent manner, only COR lengthened the period of all four reporters (Figure 2, 3, and S6). The period of TOC1 was not affected by MJ and JA-Ile and that of CCA1 was not affected by MJ.

Such a stronger role of COR in clock regulation relative to JA-Ile and MJ is consistent with previous studies that show more potent effect of COR than some JA molecules on other physiological processes [14, 27, 28]. It is possible that pathogens could use COR through a specific mechanism(s) to hyperactivate the JA signaling. Indeed, COR was shown to bind with a higher affinity to the JA receptor COI1 than plant-derived JA molecules [29, 30]. Downstream of COI1, the JA signaling is highly modular; both the JA signaling repressors (JAZ proteins) and activators (MYC proteins) belong to protein families, members of which interact with different proteins to influence multiple biological processes [31]. Therefore, it is possible that the COR-COI1 complex could selectively target some JAZ proteins for degradation, leading to a stronger or differential impact on MYC proteins and other signaling targets, such as the circadian clock. In addition to a differential perception of COR and JA molecules that could cause differences in regulating the circadian clock and other biological processes, the different efficacy between COR and other JA molecules could also be due to the solubility, uptake efficiency, stability, and catabolism of each compound in plants.

Our data further demonstrate that the four clock reporters used in this study showed different responses to COR, MJ, and JA-Ile treatments. Such a differential response of clock reporter genes to external treatments has been reported previously in response to nutrient status, ROS, phytohormones, temperature, and photoperiod [6–8, 32–35]. The differential response of these reporters may reflect tissue specific gene expression that allows differential clock response to the chemicals in separate tissues [36]. Alternatively, there may be different clocks functioning simultaneously with different rhythms in the same tissue or even in the same cell [37]. Together they support the plasticity of the circadian clock that may create flexibility for plants to respond to various external stimuli [38].

Manipulation of host circadian clock may represent a common strategy of microbes to suppress host immunity. How pathogen-produced specific molecules modulate host clock activity and defense responses still remains largely unknown. Our finding of the role of the pathogen-derived molecule COR in modulating Arabidopsis clock opens a new and exciting research direction to elucidate the molecular mechanisms underlying clock-defense interplay during host-pathogen interactions. Our data also illustrate the circadian clock being decentralized, which likely allows organisms to adapt to the changing environment in the presence of pathogens and other biotic and abiotic stresses.

## Methods

### Plant materials

All plants used in this report are in the Col-0 background. Plants were grown in growth chambers with 180 μmol m^−2^ s^−1^ photo flux density, 60% humidity, 22°C, and a 12 h light/12 h dark (LD) cycle. The clock reporter lines, *CCA1:LUC* in Col-0 or in *coi1-17* and *GRP7:LUC* in Col-0, were described previously [5]. Col-0 expressing *TOC1:LUC* or *PRR7:LUC* were kindly provided by C. R. McClung at Dartmouth College.

### RNA-seq analysis

Two sets of high-resolution RNA-seq data from 100 μM MJ or 5 μM COR treated samples were used for gene expression analysis [18, 19]. The log_2_ transformed fold changes of expression of circadian genes (Table S1) and defense genes (Table S2) were used to generate the heatmap using the heatmap.2 function in R package gplots. The circadian genes were annotated to be related to rhythmic processes according to Arabidopsis Information Resources (Table S1) and the defense genes were reported previously [21] (Table S2).

### qRT-PCR analysis

RNA extraction and qRT-PCR were performed as previously described [39, 40]. Primers used in qRT-PCR are listed in Table S3.

### Luciferase assay

Seedlings expressing the reporter gene *LUCIFERASE* (*LUC*) under the control of a clock-regulated promoter were grown on 1/2 MS media with 1% sucrose in LD and at 22°C for five days. Seedlings were transferred to 96-well plates containing 200μl of 1/2 MS medium with 0.5% sucrose, 0.4% agar, and 0.25mM D-luciferin for 1 d in LD followed by 1 d in LL with a light intensity of 180 μmol m^−2^ s^−1^. Each well contained one seedling. Seedling treatments were conducted 25 or 37 h after light onset by adding to each well 15 μl of a chemical, using COR (1 μM or 10 μM), MJ (10 μM or 100 μM), JA-Ile (10 μM or 100 μM), or sterile water as mock treatment. The dosages used for MJ, JA-Ile, and COR were chosen based on the published literature (for examples, [15, 16, 18, 19, 27, 29, 41–43]) and our preliminary experiments to test the concentrations for each chemical that induced changes of clock activity in a dosage dependent manner but did not cause overstress in plants. MJ and JA-Ile treatments with 10 μM and 100 μM demonstrated dosage-dependent phenotypes, including clock activity and seeding growth. But 100 μM coronatine drastically stunted plant growth and induced high anthocyanin production, suggesting plants under extreme stress. Thus, we used 1 and 10 μM for COR in this report.

Immediately after the treatments, the plants were measured for luminescence with an Omega Luminescence Reader (BMG LABTECH, Inc.) in LL with 90 μmol m^−2^ s^−1^ photon flux density. LUC activity was measured at 1-h intervals for five days and analyzed for amplitude, period, and phase with the R package MetaCycle [44]. All luciferase assay experiments were repeated three times with similar results.

## Acknowledgements

We thank the members in the Lu laboratory for their assistance in this work. We thank Drs. C. Robertson McClung, Gregg Howe, and Alex Webb for helpful discussions about this report. This work was partially supported by a grant from National Science Foundation (NSF 1456140) to HL.

## Author contributions

MG did clock assays, qRT-PCR, and bacterial growth. CZ did bioinformatics analysis of gene expression using RNAseq data. HL designed experiments and wrote the manuscript with help from MG and CZ.

## Additional Information

Supplementary information accompanies this paper at http://www.nature.com/srep Competing financial interests: The authors declare no competing financial interests.

## Supplemental Figure Legends

**Figure S1. COR is required for *P. syringae* virulence in LL.** (A) Bacterial growth. (B) Images to show disease symptoms. 25-d-old plants grown in LD were transferred to LL for 1 d followed by spraying with *P. syringae* 25 h after onset of LL. Statistical analysis was performed with One-way ANOVA with post-hoc Tukey HSD test (asterisks for P<0.001).

**Figure S2. Expression of selected circadian genes with MJ or COR treatment.** (A) MJ treatment. (B) COR treatment. Gene expression plots were generated by ggplot2 package in R using time-series RNA-seq data from MJ (A) or COR (B) treated samples vs. mock treated samples [18, 19].

**Figure S3. Heatmap analysis of expression of defense genes with MJ or COR treated samples**. (A) Gene expression heatmap of MJ-treated samples. (B) Gene expression heatmap of COR-treated samples. Differential gene expression values in (A) and (B) were obtained from a comparison with mock treated samples [18, 19] followed by a Log_2_ transformation.

**Figure S4. Gene expression analysis by qRT-PCR.** 7 d old LD-entrained seedlings were transferred to LL for 1 d and then treated with MJ (100 μM) or COR (10 μM) for gene expression analysis by qRT-PCR. Water was included as a mock treatment. Expression of CCA and LUX with MJ treatment was shown previously [5]. These experiments were repeated two times with similar results.

**Figure S5. Seedling growth inhibition assays.** (A) Relative seedling leaf area with COR treatment. (B) Relative seedling leaf area with MJ treatment. (C) Relative seedling leaf area with JA-Ile treatment. At the end of luminescence recording, seedlings were photographed and measured for leaf area with ImageJ. The average leaf area of water-treated samples of each genotype was set to 1 and used to calculate the relative leaf area of chemical-treated seedlings of the same genotype. Data represent mean ± SEM (n=12). Statistical analysis was performed by One-way ANOVA post-hoc Tukey HSD test. Different letters indicate significant difference among the samples treated at the same time point (P<0.05). These experiments were repeated three times with similar results.

**Figure S6. JA-Ile treatment affects clock activity** LD-entrained 5 d old seedlings were transferred to LL for 1 d and were treated with JA-Ile at 25 h (top of each panel) or 37 h (bottom of each panel). Luminescence was recorded at 1-h intervals for five days and analyzed for clock activity. A1-D1 Expression of *TOC1:LUC* in Col-0. A2-D2 Expression of *PRR7:LUC* in Col-0. A1-A2 Luminescence traces. RLU: Relative luminescence units. The color indicates JA-Ile concentration, black for 0, magenta for 10 μM, and gray for 100 μM. B1-B2 Normalized amplitude. The amplitude of the reporter was normalized to the relative leaf area shown in Figure S5. C1-C2 Period. D1-D2 Phase shift. Data represent mean ± SEM (n=12). Statistical analysis was performed by One-way ANOVA post-hoc Tukey HSD test. Different letters indicate significant difference among the samples (P<0.05). These experiments were repeated three times with similar results.

