## Supplementary material for "Coronatine is More Potent than Jasmonates in Regulating Arabidopsis Circadian Clock": Combined_Suppl_Materials.pdf

**Running title:** Coronatine regulates clock.

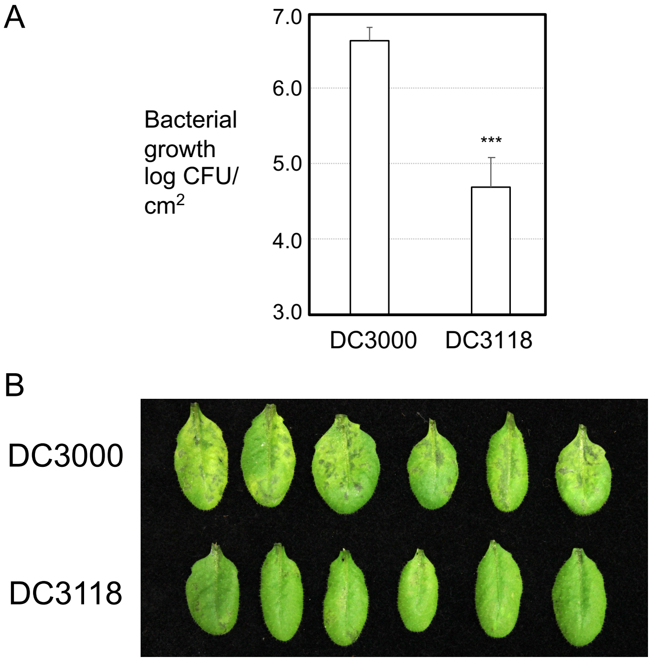
A
7.0
6.0
5.0
4.0
3.0
Bacterial growth log CFU/cm2
***
DC3000
DC3118
B
DC3000
DC3118

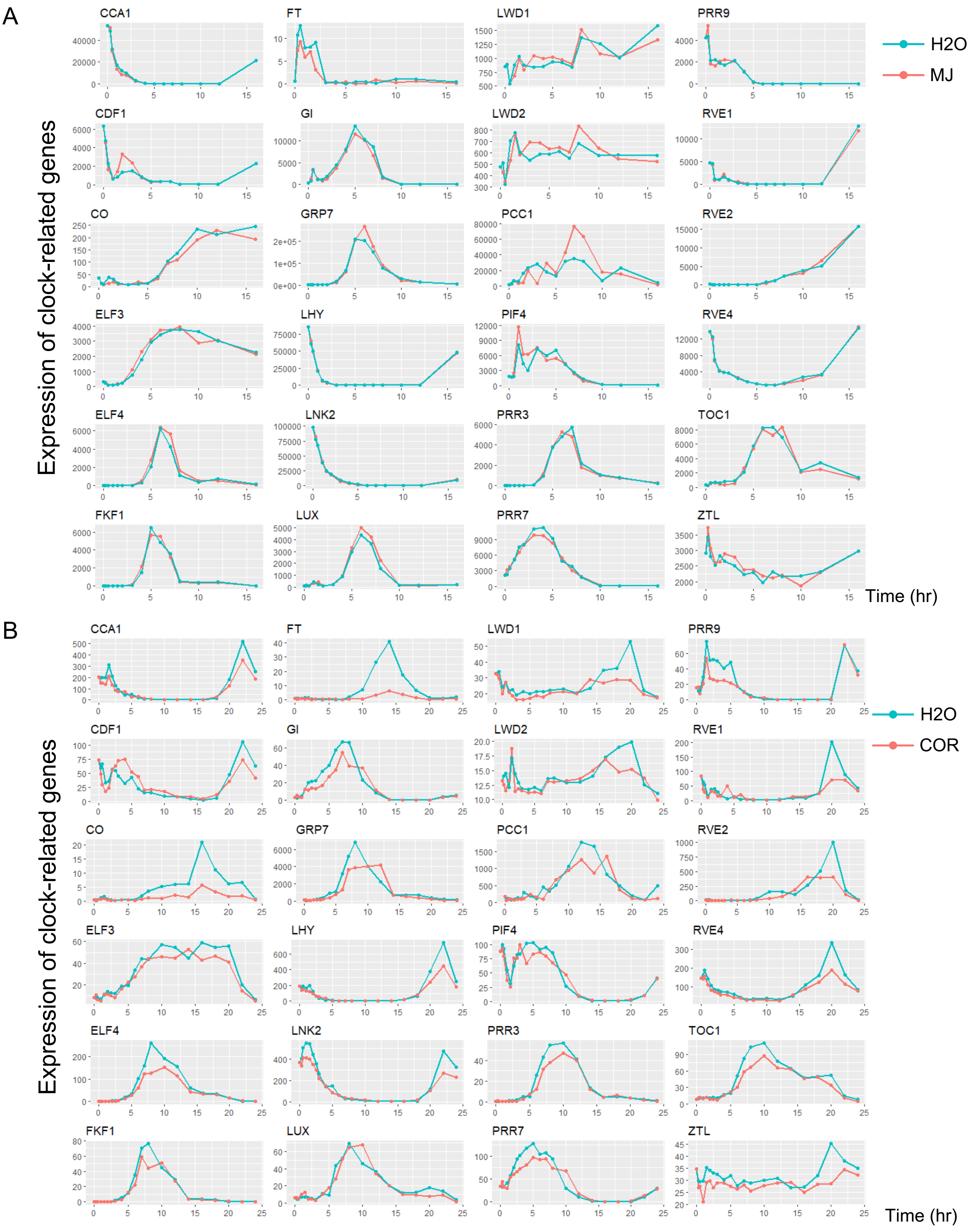
A
CCA1
FT
LWD1
PRR9
CDF1
GI
LWD2
RVE1
CO
GRP7
PCC1
RVE2
ELF3
LHY
PIF4
RVE4
ELF4
LNK2
PRR3
TOC1
FKF1
LUX
PRR7
ZTL
Expression of clock-related genes
Time (hr)
H2O
MJ
B
CCA1
FT
LWD1
PRR9
CDF1
GI
LWD2
RVE1
CO
GRP7
PCC1
RVE2
ELF3
LHY
PIF4
RVE4
ELF4
LNK2
PRR3
TOC1
FKF1
LUX
PRR7
ZTL
Expression of clock-related genes
Time (hr)
H2O
COR

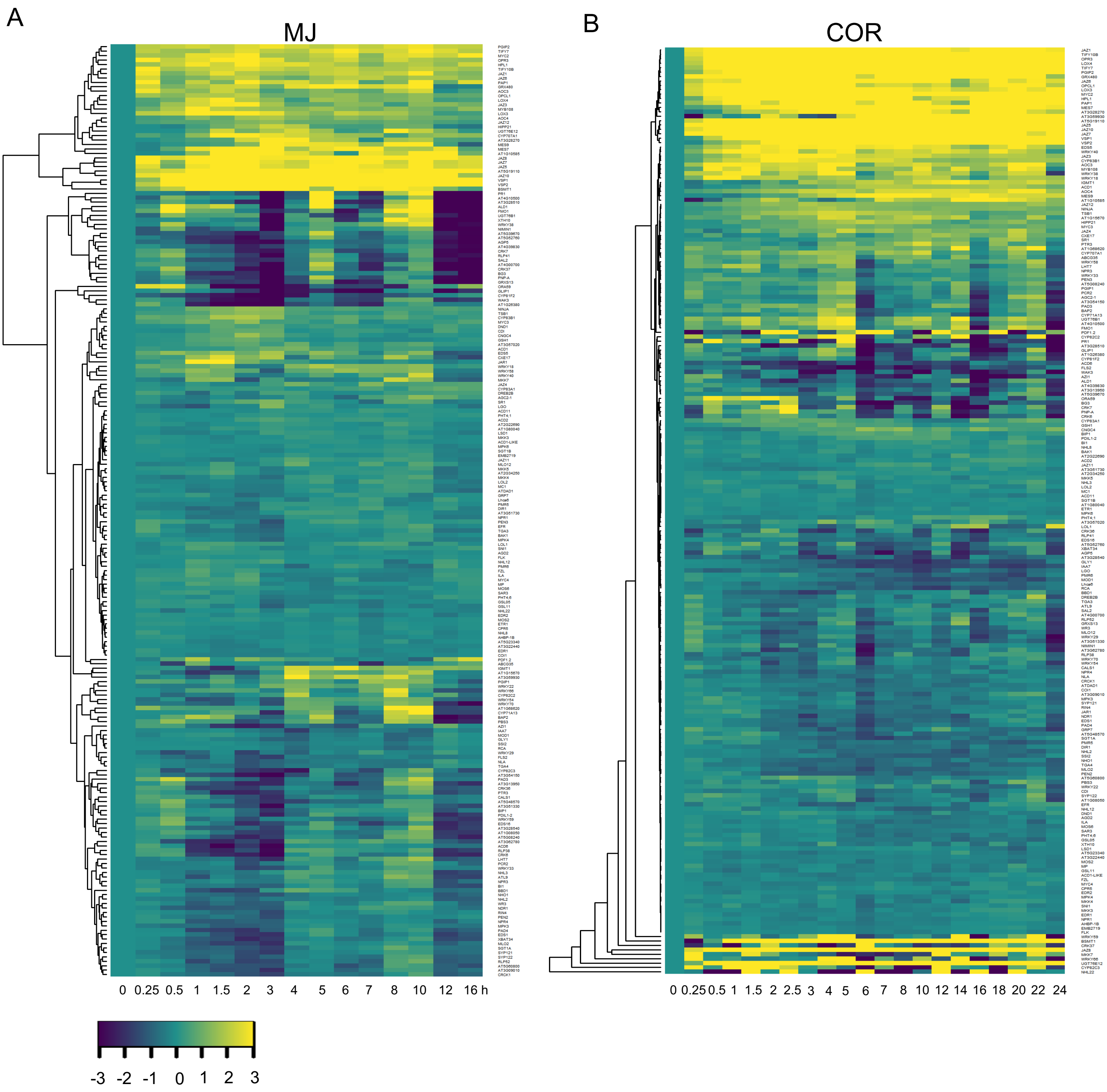
A
MJ
0 0.25 0.5 1 1.5 2 3 4 5 6 7 8 10 12 16 h
-3 -2 -1 0 1 2 3
B
COR
0 0.25 0.5 1 1.5 2 2.5 3 4 5 6 7 8 10 12 14 16 18 20 22 24

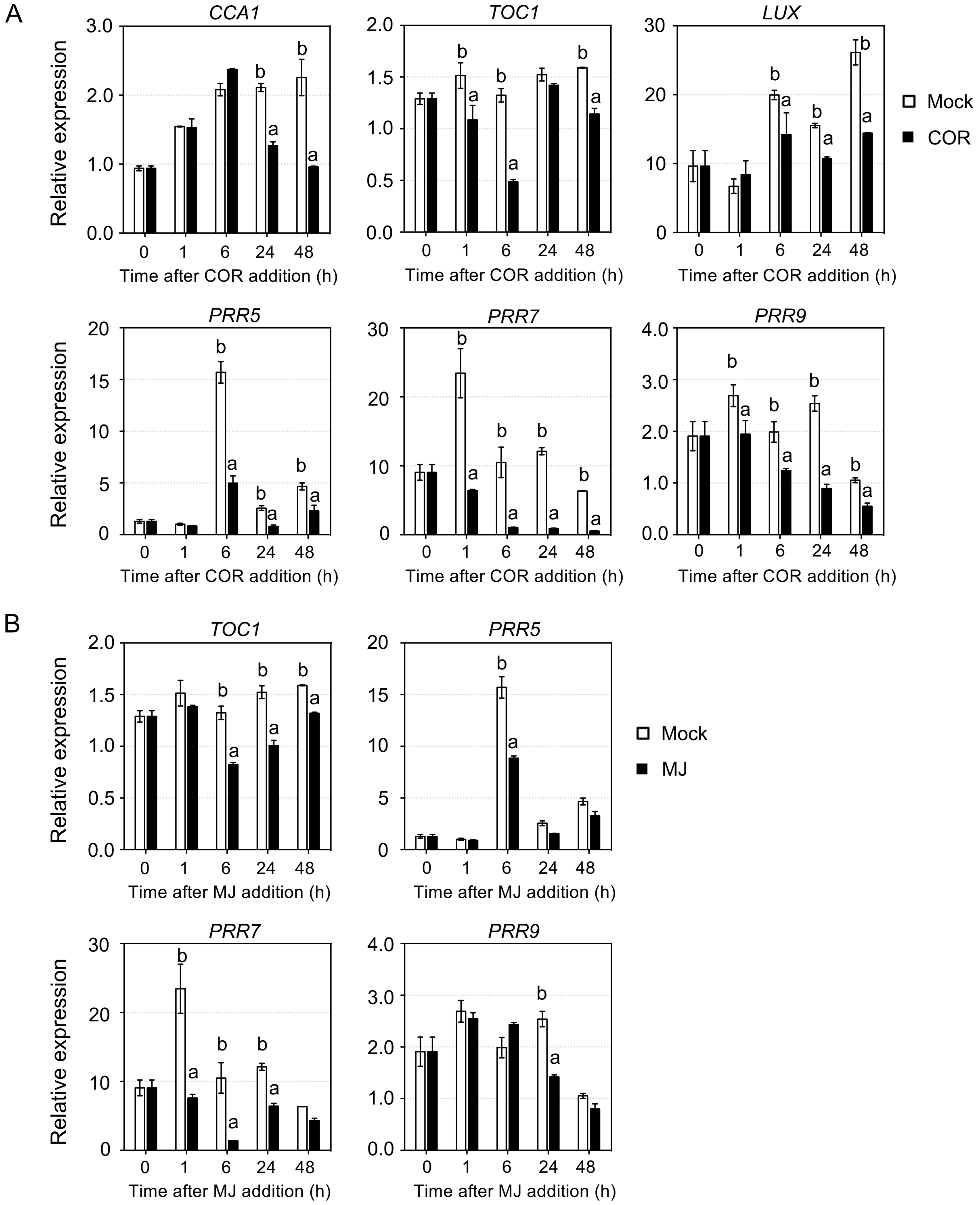
A
CCA1
TOC1
LUX
PRR5
PRR7
PRR9
Relative expression
Time after COR addition (h)
0
1
6
24
48
Mock
COR
a
b
B
TOC1
PRR5
PRR7
PRR9
Relative expression
Time after MJ addition (h)
Mock
MJ

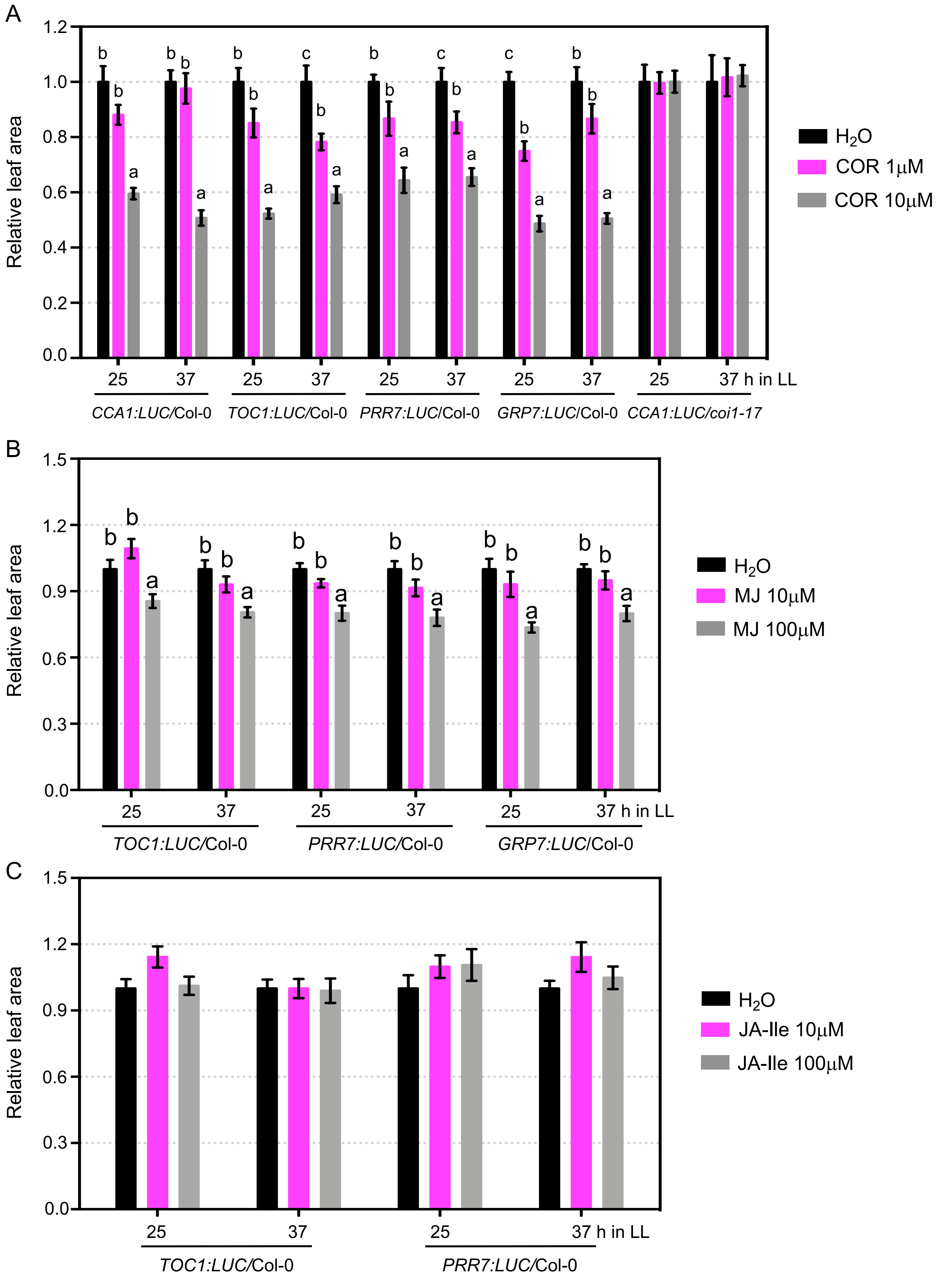
A
Relative leaf area
25
37
25
37
25
37
25
37
25
37 h in LL
CCA1:LUC/Col-0
TOC1:LUC/Col-0
PRR7:LUC/Col-0
GRP7:LUC/Col-0
CCA1:LUC/coi1-17
H2O
COR 1μM
COR 10μM
B
Relative leaf area
TOC1:LUC/Col-0
PRR7:LUC/Col-0
GRP7:LUC/Col-0
H2O
MJ 10μM
MJ 100μM
C
Relative leaf area
TOC1:LUC/Col-0
PRR7:LUC/Col-0
H2O
JA-Ile 10μM
JA-Ile 100μM

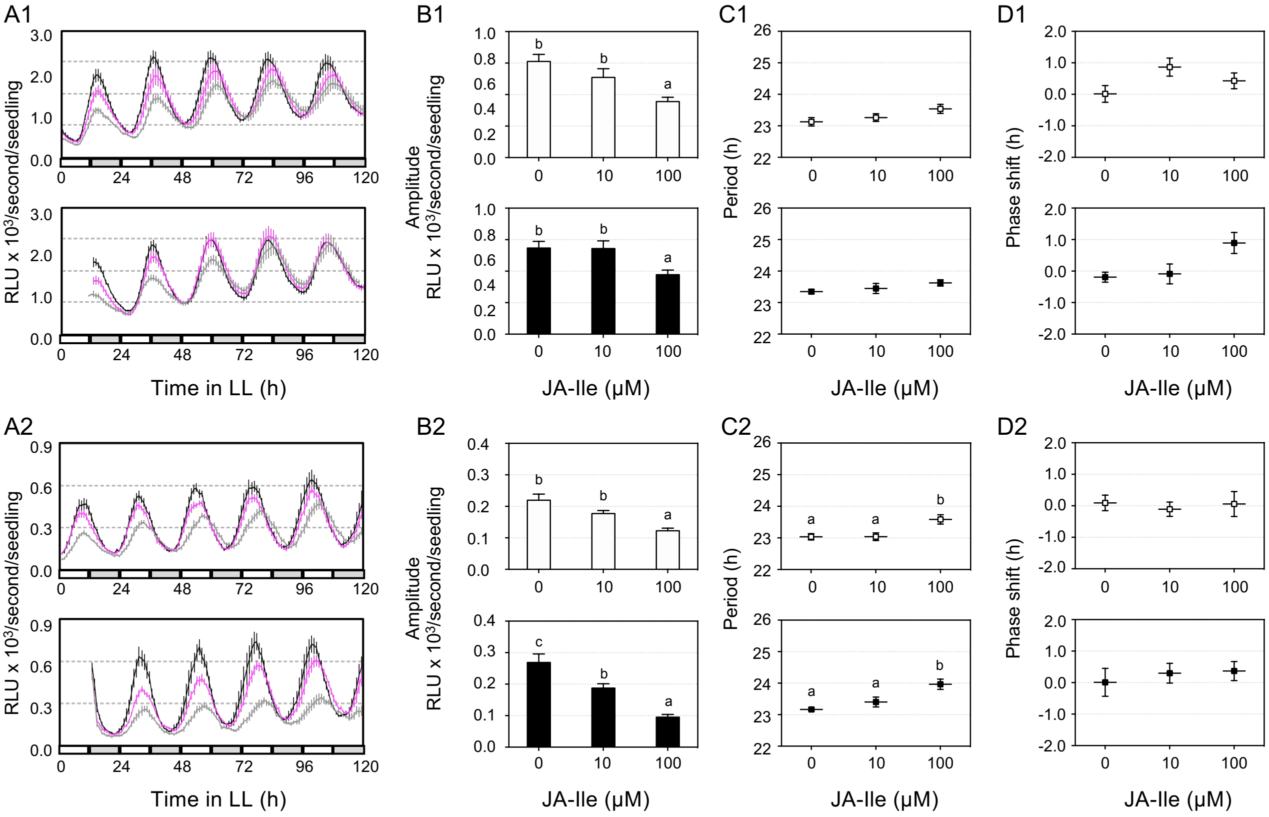
A1
B1
C1
D1
A2
B2
C2
D2
RLU x 10³/second/seedling
Time in LL (h)
Amplitude
RLU x 10³/second/seedling
JA-Ile (µM)
Period (h)
Phase shift (h)

**Table S1. Rhythmic processe-related genes used in this report.**

| AGI |
|---|
| AT1G01060 |
| AT1G04400 |
| AT1G09340 |
| AT1G09530 |
| AT1G09570 |
| AT1G10470 |
| AT1G12910 |
| AT1G15950 |
| AT1G17070 |
| AT1G17455 |
| AT1G18330 |
| AT1G22770 |
| AT1G27450 |
| AT1G31500 |
| AT1G56650 |
| AT1G59940 |
| AT1G65480 |
| AT1G68050 |
| AT1G68830 |
| AT1G68850 |
| AT1G69690 |
| AT1G72630 |
| AT1G76320 |
| AT1G77180 |
| AT1G80820 |
| AT2G06255 |
| AT2G16365 |
| AT2G17840 |
| AT2G18170 |
| AT2G18790 |
| AT2G18915 |
| AT2G21070 |
| AT2G21150 |
| AT2G21660 |
| AT2G23070 |
| AT2G23080 |
| AT2G25930 |
| AT2G29950 |
| AT2G31870 |
| AT2G32250 |
| AT2G32950 |
| AT2G35970 |
| AT2G37000 |
| AT2G37130 |
| AT2G40080 |
| AT2G41310 |
| AT2G42540 |
| AT2G43010 |
| AT2G43280 |
| AT2G44680 |
| AT2G46340 |
| AT2G46790 |
| AT2G46830 |

AT3G01060
AT3G04910
AT3G06080
AT3G06500
AT3G07500
AT3G07650
AT3G09600
AT3G11540
AT3G12320
AT3G14620
AT3G19720
AT3G20810
AT3G21890
AT3G22170
AT3G22231
AT3G22380
AT3G24050
AT3G26640
AT3G45780
AT3G46640
AT3G46780
AT3G48100
AT3G48360
AT3G50000
AT3G52180
AT3G54500
AT3G54810
AT3G55960
AT3G56480
AT3G57040
AT3G58850
AT3G59060
AT3G59470
AT3G60250
AT3G61070
AT4G00150
AT4G02570
AT4G02630
AT4G03390
AT4G04920
AT4G04970
AT4G08920
AT4G09970
AT4G10180
AT4G12850
AT4G13930
AT4G15090
AT4G15248
AT4G17640
AT4G18020
AT4G18290
AT4G19990
AT4G20370
AT4G21960
AT4G24470
AT4G25100

AT4G25470
AT4G25480
AT4G25490
AT4G26150
AT4G26700
AT4G28610
AT4G30200
AT4G30350
AT4G31120
AT4G32890
AT4G34680
AT4G36240
AT4G36930
AT4G37520
AT4G37930
AT4G38960
AT5G02120
AT5G02810
AT5G02840
AT5G05660
AT5G08330
AT5G10140
AT5G11260
AT5G12050
AT5G13930
AT5G15840
AT5G15850
AT5G17300
AT5G23730
AT5G24470
AT5G25830
AT5G37260
AT5G47080
AT5G48890
AT5G51810
AT5G51990
AT5G52250
AT5G52310
AT5G52660
AT5G52910
AT5G56860
AT5G57360
AT5G58140
AT5G59560
AT5G59570
AT5G60100
AT5G61380
AT5G62430
AT5G63860
AT5G64120
AT5G64170
AT5G64813
AT5G67380

**Table S2. Defense-related genes used in this report.**

| AGI |
|---|
| AT1G02170 |
| AT1G02450 |
| AT1G02860 |
| AT1G03160 |
| AT1G03850 |
| AT1G05570 |
| AT1G06160 |
| AT1G08050 |
| AT1G08720 |
| AT1G10585 |
| AT1G11310 |
| AT1G13280 |
| AT1G14870 |
| AT1G15210 |
| AT1G15670 |
| AT1G17380 |
| AT1G17420 |
| AT1G18350 |
| AT1G19150 |
| AT1G19180 |
| AT1G19250 |
| AT1G19850 |
| AT1G20200 |
| AT1G20510 |
| AT1G21100 |
| AT1G21240 |
| AT1G22070 |
| AT1G26380 |
| AT1G28480 |
| AT1G30135 |
| AT1G32210 |
| AT1G32340 |
| AT1G32540 |
| AT1G32640 |
| AT1G33520 |
| AT1G35230 |
| AT1G48500 |
| AT1G51660 |
| AT1G56650 |
| AT1G59870 |
| AT1G64280 |
| AT1G64790 |
| AT1G64980 |
| AT1G66340 |
| AT1G68620 |
| AT1G70700 |
| AT1G72450 |
| AT1G72520 |
| AT1G74710 |
| AT1G74950 |
| AT1G75380 |
| AT1G77510 |
| AT1G80040 |

AT1G80460
AT1G80590
AT1G80680
AT1G80840
AT2G05990
AT2G06050
AT2G13810
AT2G14610
AT2G14620
AT2G18660
AT2G21660
AT2G21900
AT2G22690
AT2G23560
AT2G29650
AT2G30770
AT2G34250
AT2G34600
AT2G34690
AT2G35000
AT2G35960
AT2G38470
AT2G39200
AT2G39730
AT2G39940
AT2G40690
AT2G40750
AT2G43710
AT2G43790
AT2G44490
AT2G45760
AT2G46370
AT3G01080
AT3G04610
AT3G06490
AT3G09010
AT3G10525
AT3G11020
AT3G11340
AT3G11480
AT3G11650
AT3G11820
AT3G13950
AT3G17860
AT3G20600
AT3G21220
AT3G22440
AT3G23050
AT3G23120
AT3G25010
AT3G25070
AT3G25250
AT3G25780
AT3G26830
AT3G28270
AT3G28510

AT3G28540
AT3G43440
AT3G44880
AT3G45640
AT3G46660
AT3G48090
AT3G51330
AT3G51730
AT3G52400
AT3G52430
AT3G54150
AT3G54920
AT3G56400
AT3G57020
AT3G57240
AT3G59100
AT3G59930
AT3G62780
AT4G00700
AT4G01250
AT4G01370
AT4G02150
AT4G03550
AT4G04490
AT4G04500
AT4G09590
AT4G10500
AT4G11260
AT4G12470
AT4G13770
AT4G14365
AT4G14400
AT4G15440
AT4G17880
AT4G18470
AT4G19040
AT4G19230
AT4G19660
AT4G20380
AT4G21610
AT4G23100
AT4G23140
AT4G23150
AT4G23550
AT4G23570
AT4G25650
AT4G28910
AT4G31500
AT4G31800
AT4G31950
AT4G31970
AT4G33430
AT4G33680
AT4G35180
AT4G37000
AT4G37150

AT4G39030
AT4G39830
AT5G01820
AT5G06320
AT5G06860
AT5G06870
AT5G06950
AT5G08240
AT5G10030
AT5G13220
AT5G13320
AT5G15410
AT5G16080
AT5G17450
AT5G19110
AT5G20480
AT5G20900
AT5G22570
AT5G23340
AT5G24770
AT5G24780
AT5G25910
AT5G28540
AT5G39670
AT5G40440
AT5G40990
AT5G44370
AT5G44420
AT5G45110
AT5G46050
AT5G46330
AT5G46760
AT5G47120
AT5G48485
AT5G48570
AT5G50200
AT5G52760
AT5G54250
AT5G54810
AT5G57220
AT5G58600
AT5G58940
AT5G60800
AT5G64000
AT5G64930

**Table S3. qRT-PCR primers used in this report.**

| Gene Name | Primer name | Primer sequence |
|---|---|---|
| CCA1 | CCA1-qPCR-f | TCGAAAGACGGGAAGTGGAACG |
| | CCA1-qPCR-r | GTCGATCTTCATTGGCCATCTCAG |
| *LUX* | LUX-qPCR-f | AACACCTGTTCCTCCACAGAGC |
| | LUX-qPCR-r | TCCAACATTACCGCTGCTACCG |
| *PRR5* | PRR5-qPCR-f | AGCTTTCACACGGTACGTTCAC |
| | PRR5-qPCR-r | TTGGAGGCGGTTCAGATGTATTG |
| *PRR7* | PRR7-qPCR-f | AAGCGGAAGTGGAAGTGGTAGC |
| | PRR7-qPCR-r | TCCGGCTTTGGTATCGTACCTTC |
| PRR9 | PRR9-qPCR-f | TGAAGTGTATGCTGAGAGGTGCT |
| | PRR9-qPCR-r | AGCAGTAGGATCATCACGCAAAG |
| TOC1 | TOC1-qPCR-f | TTAGGTCCACCAACCCACAGAGA |
| | TOC1-qPCR-r | AGGAGCAGTAGCAACAGACCACT |
